# Energetic stress and infection generate immunity-fecundity tradeoffs in *Drosophila*

**DOI:** 10.1101/318568

**Authors:** Justin L. Buchanan, Colin D. Meiklejohn, Kristi L. Montooth

## Abstract

Physiological responses to short-term environmental stress, such as infection, can have long-term consequences for fitness, particularly if the responses are inappropriate or nutrient resources are limited. Genetic variation affecting energy acquisition, storage, and usage can limit cellular energy availability and may influence resource-allocation tradeoffs even when environmental nutrients are plentiful. Here, we utilize well-characterized *Drosophila* mitochondrial-nuclear genotypes to test whether disrupted energy metabolism interferes with nutrient-sensing pathways, and whether this disruption has consequences for tradeoffs between immunity and fecundity. We find that this energetically compromised genotype is resistant to rapamycin – a drug that stimulates nutrient-sensing pathways that are activated when resources are limited. Resource limitation also compromises survival in energetically-compromised genotypes, suggesting that this genotype may have little excess energetic capacity and fewer cellular nutrients, even when environmental nutrients are not limiting. Accordingly, we find that immune function is compromised in this genotype, but only in females, and that these females experience immunity-fecundity tradeoffs that are not evident in genotypic controls with normal energy metabolism. Thus, genetic variation in energy metabolism may act to limit the resources available for allocation to life-history traits in ways that generate tradeoffs even when environmental resources are not limiting.

## Introduction

The energy available to heterotrophic organisms is often determined by nutrients in the environment, and the dynamic allocation of these resources within the lifespan of an individual impacts life-history tradeoffs between organismal maintenance and reproduction. Nutritional stress may be caused by the lack of a single nutrient (Bergland et al. 2008; Jensen et al. 2015), improper nutrient ratios (Skorupa et al. 2008), or reduced overall food availability leading to a decrease in overall calorie consumption (Itskov et al. 2014; Ro et al. 2014). Energetic costs associated with infection are predicted to have a significant impact on survivorship and future reproduction via the allocation of limited resources between reproduction and immunity (Lochmiller and Deerenberg 2000; Harshman and Zera 2007; Schwenke et al. 2016). Energetic costs of infection can be associated with the mechanisms of disease resistance (e.g., constitutive and induced immune responses) and tolerance (Rauw 2012), reduced nutrient uptake during infection (Bonfini et al. 2016), or pathogen consumption of resources (Cressler et al. 2014; Kurze et al. 2016).

Despite the prediction that fighting infection will generate a trade-off with future reproduction, the relationship between infection and reproduction is complex. Under some conditions, adult infection decreases fecundity and the expression of reproduction genes (Short and Lazzaro 2013). However, constitutive immune expression does not always generate life-history tradeoffs (Fellous and Lazzaro 2011), and infection can even increase fecundity (Adamo 1999) and offspring quality (Stahlschmidt et al. 2013; Reavey et al. 2015). Post-infection increases in fecundity may occur if the costs of clearing infection are high enough to switch energetic resources towards short-term investment in reproduction (Cressler et al. 2015). Thus understanding how host metabolism can impact resource allocation and influence life-history tradeoffs remains an important area of research.

Investigating how genetic variation in host metabolism interacts with immunity and diet to influence life-history outcomes during periods of environmental stress (e.g., infection) is critical for understanding how immunity-fecundity tradeoffs evolve. Genetic variation affecting energy metabolism may limit the availability of cellular energy (e.g. energy stores or ATP) and influence resource-allocation tradeoffs even when environmental nutrients are not limiting. One regulatory mechanism that integrates information both from external (e.g. food availability) and internal (e.g. cellular ATP availability) inputs to impact life-history traits is the target of rapamycin (TOR) signaling pathway (Oldham and Hafen 2003). When external and internal nutrient levels are sufficient, TOR upregulates downstream target genes to promote protein synthesis. Conversely, poor nutrient levels or treatment with the drug rapamycin decreases protein production and increases recycling of cellular components via autophagy (Zheng et al. 1995; Hahn-Windgassen et al. 2005) (Fig. 1). Consistent with these effects, treating *Drosophila* with rapamycin delays development, decreases fecundity, and increases lifespan (Bjedov et al. 2010).

**Figure 1.**
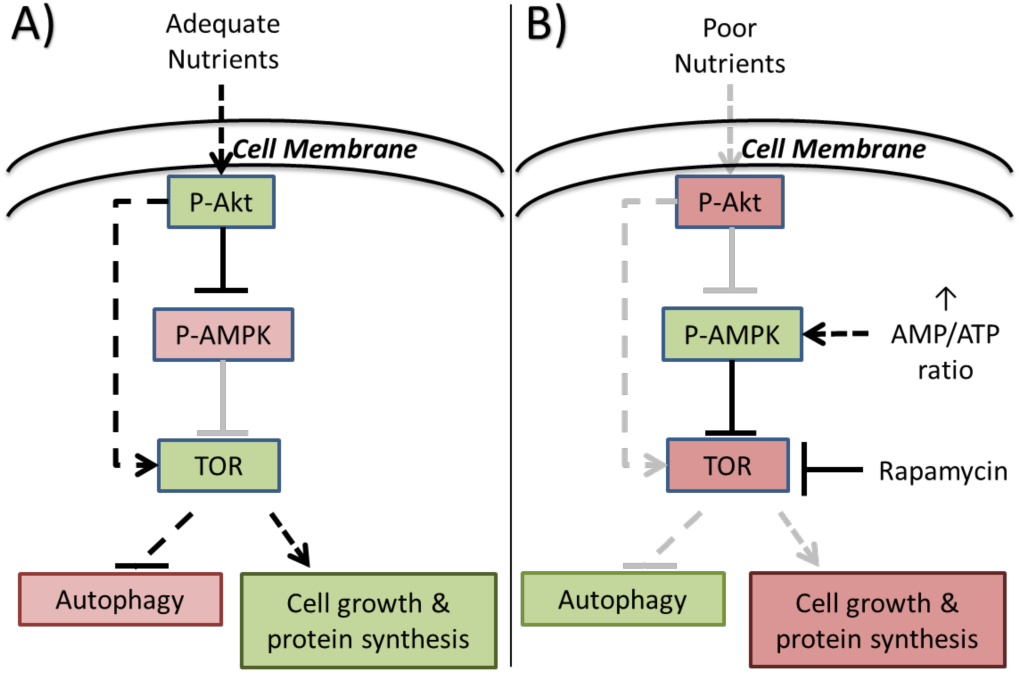
The protein Target of Rapamycin (TOR) integrates nutrient responses to regulate growth and development. A) In the presence of adequate nutrients, TOR is active, which represses autophagy and promotes cell growth. B) When nutrients are sensed as being limited either via insulin signaling, an increased AMP/ATP ratio, or artificially by exposure to the drug rapamycin, TOR is inactivated which promotes autophagy and inhibits cell growth.

To investigate how genetic variation in energy metabolism and, specifically, in mitochondrial function may mediate immune function and immunity-fecundity tradeoffs, we utilized a mitochondrial-nuclear (mito-nuclear) hybrid genotype (Montooth et al. 2010) that compromises mitochondrial oxidative phosphorylation. Compromised OXPHOS in this genotype is caused by an incompatible interaction between a single nucleotide polymorphism in the mitochondrial-encoded mt-tRNA^Tyr^ and an amino acid polymorphism in the nuclear-encoded mt-tyrosyl-tRNA synthetase that aminoacylates this mt-tRNA (Meiklejohn et al. 2013). Together, these mutations decrease female fecundity, delay development, and disrupt larval metabolic rate, indicative of inefficient energy metabolism (Hoekstra et al. 2013, 2018;Meiklejohn et al. 2013). Here we measure life-history traits in mito-nuclear genotypes under nutrient and pathogen stress to test whether genetic variation that compromises energy metabolism can limit available cellular resources and generate tradeoffs between immunity and fecundity.

## Methods

### Drosophila Genotypes and Rearing Conditions

We employed a panel of six mito-nuclear genotypes that combine mtDNAs from *D. simulans* – (*simw^501^*) and (*sm21*) – and *D. melanogaster* (*ore*) with two wild-type *D. melanogaster* nuclear genomes – *OreR* and *Aut* (Montooth et al. 2010). Of these six genotypes, only the (*simw^501^*);*OreR* mito-nuclear combination generates an incompatible interaction that decreases oxidative phosphorylation; the other five genotypes serve as wild type controls. All genotypes are maintained on our standard laboratory food at 25 °C with a 12h:12h, light:dark cycle. Three non-isocaloric food types were used during experiments: our standard laboratory food, also denoted as high yeast in this paper, (0.88% agar, 8.33% Torula Yeast, 10% Cornmeal, 0.33% Tegosept W/V and 4.66% Molasses, 1.66% 95% ethanol, 0.66% propionic acid V/V dH_2_O), low-yeast food (our standard food with 0.5% Torula Yeast W/V), and a medium-mixed diet (0.93% agar, 2.94% SAF Yeast, 6.12% Cornmeal, 12.94% sugar, 0.28% Tegosept W/V and 1.08% 95% ethanol, 0.71% propionic acid V/V dH_2_O).

### Rapamycin and diet effects on development

The first experiment was performed using all 6 genotypes and 3 rapamycin concentrations (0 μM, 2 μM and 10 μM) on medium mixed diet. 50 females and 30 males were mated for 24 hours and placed onto grape agar plates for collecting cohorts of 75 eggs every 24 hours (50 g bacto-agar, 30 mL tegosept in 10% ethanol, 500 mL grape juice, 1500 mL distilled H_2_O). A total of 5 vials of 75 eggs per genotype and rapamycin level were monitored twice a day to measure the date to first pupation, the development time of each individual, and the number of males and females that eclosed as a measure of sex-specific survival. Our estimate of sex-specific survival assumed a 50:50 sex ratio in the eggs or larvae (see below) placed in each vial.

In order to examine more concentrations of rapamycin on a high-yeast diet, genotypes with the (*sm21*) mtDNA – which did not behave differently from the (*ore*) mtDNA in the preliminary experiment – were not included in a second experiment. In this experiment, four genotypes were reared on a high-yeast food for many generations before being reared on food containing 0 μM, 5 μM, 10 μM, or 15 μM rapamycin. Males and females of each genotype were mated, and females were allowed to lay eggs for 12 hours on grape agar plates and were then removed. 50 first-instar larvae of each genotype were collected 24 hours after the egg lay by floating the larvae in 20% sucrose and 1X phosphate-buffered saline (PBS). We set up 7 replicate vials of each genotype at each rapamycin concentration, with an additional replicate vial for (*simw^501^*);*OreR* due to the increased probability that vials are non-productive with this genotype because of its overall low-fecundity and embryonic defects (Meiklejohn et al. 2013; Zhang et al. 2017). Vials were scored for development time and survival as described above.

In order to test whether control genotypes exposed to a low-yeast environment would show a decreased responsiveness to rapamycin, similar to (*simw^501^*);*OreR* on a high-yeast diet, we developed the full panel of six mito-nuclear genotypes from larvae to adult on either a high-yeast or low-yeast diet, supplemented with three levels of rapamycin (0 μM, 5 μM, and 10 μM). Males and females of each genotype were mated, and females were allowed to lay eggs for 4 hours on high or low yeast plates, respectively, and were then removed. 100 first-instar larvae of each genotype were collected 30 hours after the egg lay by floating the larvae in 20% sucrose and 1X PBS. We set up 5 replicate vials of each genotype, yeast, and rapamycin combination. Vials were scored for development time and survival as described above.

### Bacterial Infection

To test whether compromised energy metabolism decreases the ability to survive bacterial pathogen infection, we infected virgin 1-day old adults of all six mito-nuclear genotypes with the natural pathogen *Providencia rettgeri* (Juneja and Lazzaro 2009; Short and Lazzaro 2013). Individuals were either sham infected with 1X PBS or infected with *P. rettgeri* in 1X PBS at a concentration of 1.0 O.D. using a 0.1 mm needle (TedPella 13561-50). This infection protocol results in moderate lethality, with infection stabilizing by day 4 (Sackton et al. 2010; Howick and Lazzaro 2014; Duneau, Ferdy, et al. 2017). Flies were then placed in groups of 30 males or females on standard food and counted twice daily for survival for 10 days. After 5 days, individuals that were still alive were transferred to fresh food. Five replicate groups were measured for survival for each genotype, sex, and infection treatment (sham versus pathogen) combination. In a parallel infection setup, fecundity was measured using 15-20 females of each genotype/treatment that had survived to 5 days post infection. These females were mated with wild-type males from an outbred *D. melanogaster* population that is genetically distinct from the nuclear backgrounds of the focal genotypes. Mated females were allowed to lay eggs for 72 hours, transferring both males and females to a new vial every 24 hours.

### Statistical Analyses

Development time to adult eclosion was analyzed using a linear mixed-effects model with mtDNA, nuclear genotype, sex, treatment (rapamycin, yeast, and infection) and their interactions as fixed effects, and vial as a random variable. Rapamycin concentration was treated as a continuous variable. Tukey tests were performed with Holm correction to look at significance due to individual genotype and/or treatment effects. The same fixed effects were used included in a generalized linear model analysis of survivorship, but with a binomial distribution for the survival data. For models that include sex as a factor, the assumption was made that an equal number of male and female larvae were initially collected. No random effect of vial was required for survival analysis as each replicate vial provides one measure of the number of larvae that survived to adults. Fecundity was analyzed using linear models that included the fixed effects of day, genotype, and treatment. Due to the high variance of fecundity, outliers were removed via Grubbs test. However, analysis with and without outlier data did not produce qualitatively different results. All analyses were carried out in R version 3.1.3 (R Core Team 2015).

## Results

### Individuals with compromised energy metabolism are largely resistant to the effects rapamycin

Previous work has shown that the mito-nuclear genotype (*simw^501^*);*OreR* decreases oxidative phosphorylation (OXPHOS) activity with deleterious effects on metabolic rate, development, and female fecundity that are sensitive to energy demand (Hoekstra et al. 2013; Meiklejohn et al. 2013; Holmbeck et al. 2015; Hoekstra et al. 2018; Zhang et al. 2017). We tested whether (*simw^501^*);*OreR* flies have altered nutrient sensing even when reared on a non-limiting diet due to their predicted low levels of cellular energy. We raised this genotype and genotypic controls that have normal energy metabolism on a high-nutrient diet supplemented with increasing concentrations of rapamycin. The target of rapamycin protein (TOR) is an energy-sensing protein downstream of both the insulin receptor and AMP-activated protein kinase (AMPK) – a central regulator of cellular metabolism that responds to the abundance of AMP relative to ATP. Thus, TOR integrates multiple signals of nutrient availability and energetic status (Fig. 1).

In *Drosophila*, rapamycin extends development in a dose-dependent manner (Zhang et al. 2000; Wang et al. 2016). We found that (*simw^501^*);*OreR* was remarkably resistant to the effects of rapamycin on both development time and survival to adulthood (Fig. 2). We found a significant interaction between mtDNA genotype, nuclear genotype, and rapamycin concentration on development time (mtDNA*nuclear*rapamycin: *F*_1,83_ = 9.49, *P* = 0.0028) that was attributable to an attenuated response of (*simw^501^*);*OreR* development time to rapamycin treatment, compared to the other five genotypes (Fig. 2A). (*simw^501^*);*OreR* had the longest development time in the absence of rapamycin (*P* < 1.75e-09 for all Tukey contrasts) but developed faster than all other genotypes at all concentrations of rapamycin; this difference was statistically significant on 10μM rapamycin (*P* < 5.27e-06 for all Tukey contrasts). There was a nearly significant four-way interaction between mitochondrial genotype, nuclear genotype, rapamycin, and sex on development time (mtDNA*nuclear*rapamycin*sex: *F*_1,1850_ = 3.73, *P* = 0.054) caused by a somewhat dampened effect of rapamycin in extending male relative to female development time in some genotypes (Supplementary Fig. 1 A&B).

**Figure 2.**
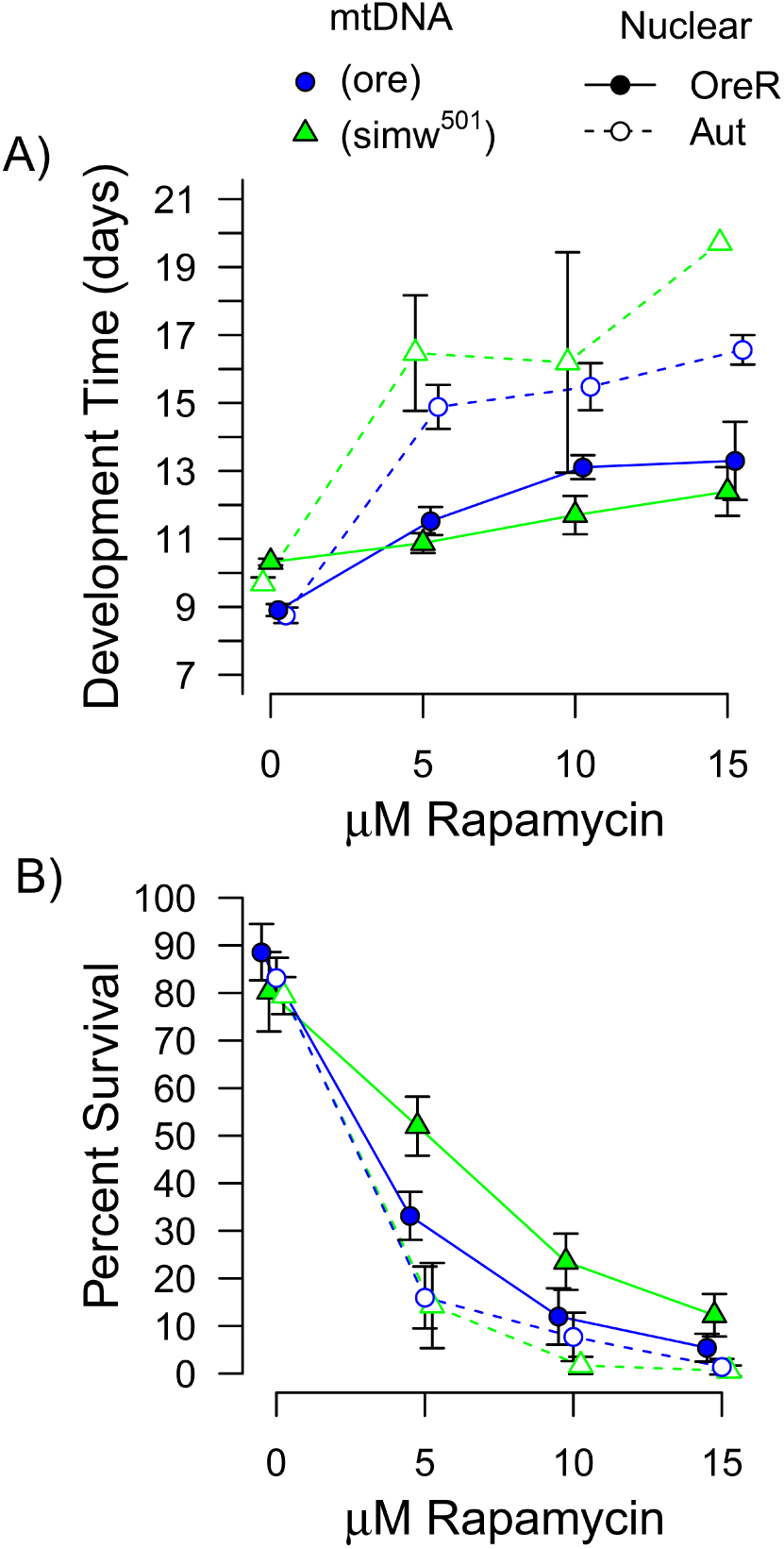
An energetically compromised genotype (*simw^501^*);*OreR* was resistant to the drug rapamycin. A) The effect of rapamycin to increase development time was attenuated in (*simw^501^);OreR* relative to control genotypes. B) (*simw^501^*);*OreR* had similar survival to genetic controls in the absence of rapamycin, but had the highest survival in the presence of rapamycin. Points are average trait values across 7-8 replicate vials with 95% CI. Low survivorship of the *Aut* nuclear background accounts for the increase in variance and lack of error bars at the highest doses of rapamycin. Female and male data are combined in this figure, as the patterns were similar between the sexes (Supplementary Fig. 1).

In addition to delaying development, rapamycin caused significant concentration-dependent mortality in all genotypes (Fig. 2B). However, this effect was also significantly attenuated in (*simw^501^*);*OreR* relative to the control genotypes (mtDNA*nuclear*rapamycin: *χ^2^*_1,220_ = 28.24, *P* < 0.0001), a pattern that was independent of sex (mtDNA*nuclear*rapamycin*sex: *χ^2^*_1,216_ = 0.04, *P* = 0.8) (Fig. 2B, Supplemental Fig. 1 C&D). In fact, when first-instar larvae were developed on food containing rapamycin, (*simw^501^*);*OreR* showed the highest larval-to-adult survival of all genotypes (Fig. 2B). In the absence of rapamycin, (*simw^501^*);*OreR* had wild-type larval-to-adult survival (mtDNA*nuclear: *χ^2^*_1,54_ = 2.56, *P* = 0.11). In contrast, when eggs, rather than larvae, were transferred to food containing no rapamycin, (*simw^501^*);*OreR* showed significantly lower egg-to-adult survival, relative to other genotypes (mtDNA*nuclear: *χ^2^*_2,53_ = 17.8, *P* < 0.001) (Supplementary Fig. 2), consistent with prior work indicating that compromised energy metabolism has a strong negative effect on maternal provisioning of eggs that compromises embryogenesis in this genotype (Hoekstra et al. 2013; Zhang et al. 2017). In summary, (*simw^501^*);*OreR* individuals appear to be relatively resistant to the effects of rapamycin for survival to adulthood, in addition to development time, suggesting that this genotype may have less responsive TOR signaling as a consequence of a deficient cellular energetic state even when provided a high-nutrient diet.

### Developmental effects of decreased dietary nutrients are genotype and sex specific

Prior studies have demonstrated that dietary yeast affects *Drosophila* development and ovary size (Bergland et al. 2008; Becher et al. 2012). Yeast are thought to be an important source of dietary amino acids for *Drosophila*, and limiting dietary amino acids slows *Drosophila* development, possibly via TOR (Colombani et al. 2003; Oldham and Hafen 2003). We reared the mito-nuclear genotypes on a low-yeast diet across a range of rapamycin concentrations to test two hypotheses. First, we tested whether (*simw^501^*);*OreR* individuals were relatively resistant to the effects of decreased dietary yeast. Second, we aimed to test whether genotypic controls with decreased dietary nutrients became more resistant to rapamycin, analagous to (*simw^501^*);*OreR* individuals fed a non-limiting diet. While a low-yeast diet extended development in all genotypes in the absence of rapamycin, the effect was smaller for (*simw^501^*);*OreR* flies (Fig. 3 A&B). In a high-yeast environment, the development time of this genotype was nearly two days delayed relative to genotypic controls (Tukey’s *P_females_* < 0.05, *P_males_* < 0.01) – a pattern also observed on the mixed diet that was intermediate in yeast content (Tukey’s *P* < 0.001) (Supplementary Fig. 3B). However, in a low-yeast environment the developmental time of (*simw^501^*);*OreR* flies was not significantly different from genotypic controls (all Tukey’s *P* > 0.35). This inability to extend development appeared to come at a cost to female survivorship; female (*simw^501^*);*OreR* larval-to-adult survivorship was significantly reduced to 50% on a low-yeast diet, relative to control genotypes (Tukey’s *P* < 0.001) (Fig. 3C), while males had high survivorship similar to other nuclear genotypic controls under both diets (Tukey’s *P_High_* > 0.05, *P_Low_* > 0.05) (Fig. 3D).

**Figure 3.**
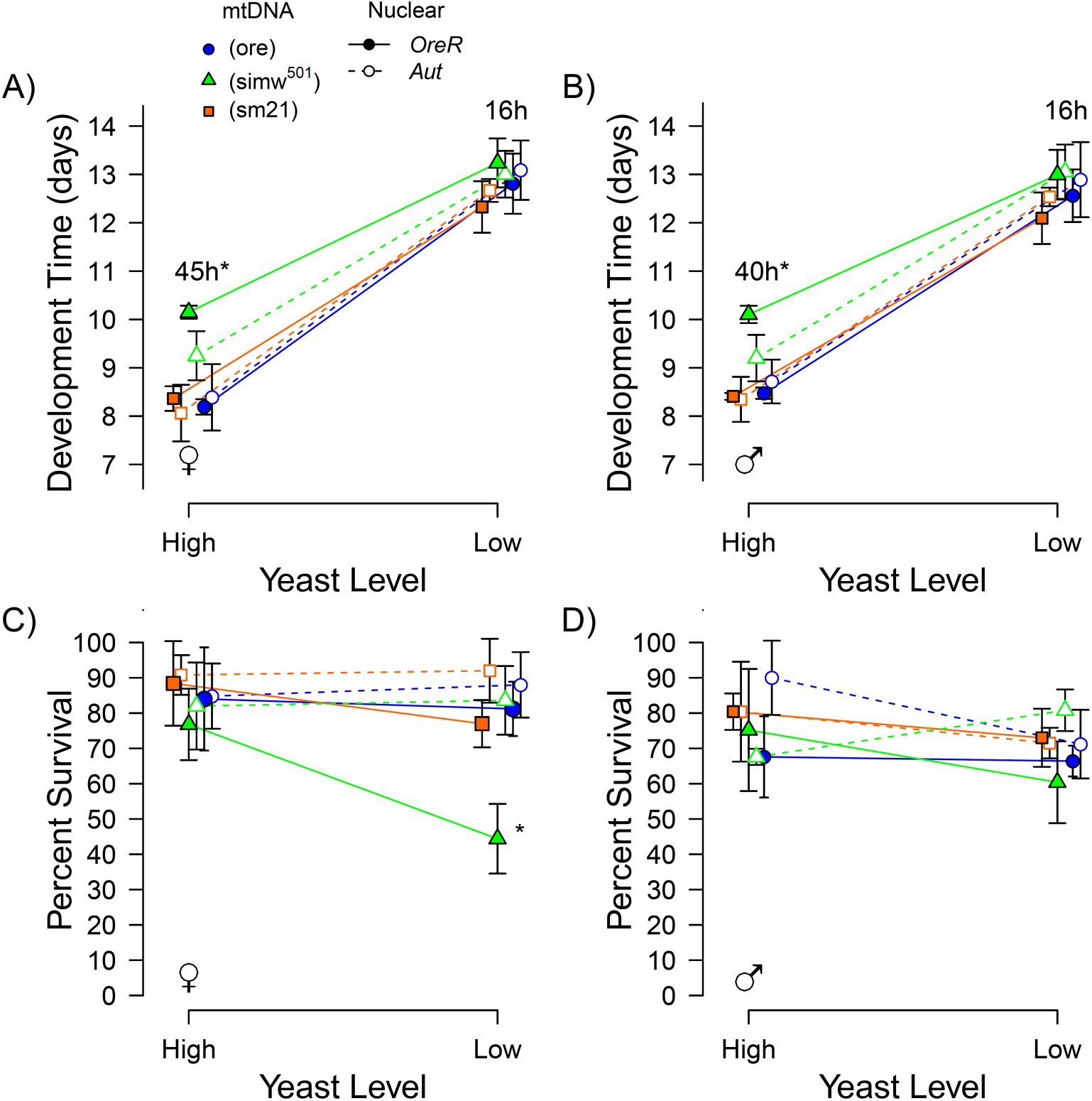
Dietary yeast modified the effects of a mitochondrial-nuclear incompatibility on development and female larval-to-adult survival. A & B) Decreased dietary yeast delayed development of all genotypes, but the response of (*simw^501^*);*OreR* to dietary yeast was less than that of control genotypes. Development of this genotype was significantly delayed relative to nuclear genotypic controls when developed on a high-yeast diet, but not when developed on a low-yeast diet. Numbers over the points indicate the difference in average development time in hours between (*simw^501^*);*OreR* and *OreR* nuclear genotypic controls. C & D) (*simw^501^*);*OreR* females had significantly decreased survival relative to all other genotypes when developed specifically on a low-yeast diet, an effect that was not observed in males. Points are average trait values across five replicate vials with 95% CI.

Flies with the *Aut* nuclear background had very low survivorship when developed on rapamycin, independent of mtDNA. This effect was enhanced on the low-yeast diet, with very few individuals surviving after greatly extended development in the presence of rapamycin. At 10 μM rapamycin on a low-yeast diet, too few flies of all genotypes survived to provide good estimates of development time (Supplementary Fig. 4). However, we were able to use two compatible mito-nuclear genotypes with the *OreR* nuclear background – (*ore*);*OreR* and (*sm21*);*OreR* – to test the prediction that control genotypes fed a low-yeast diet would be less responsive to 5 uM rapamycin, analogous to the (*simw501*);*OreR* genotype on a high-yeast diet. (*ore*);*OreR* flies developed on a low-yeast diet had a decreased response of development time to 5 μM rapamycin, relative to (*ore*);*OreR* flies developed on a high-yeast diet (Fig. 4), an effect that was somewhat stronger in females (females, yeast*rapamycin: *F* _2,46_ = 12.3, *P* < 0.001; males, yeast*rapamycin: *F* _2,40_ = 3.0, *P* = 0.059) (Supplemental Table 1). However, this pattern was not observed in (*sm21*);*OreR* (females, yeast*rapamycin: *F* _2,41_ = 0.98, *P* = 0.38; males, yeast*rapamycin: *F* _2,40_ = 0.07, *P* = 0.93) (Fig. 4). Together, our results indicate that nutrient limitation – either in the diet or by mutations affecting energy metabolism – attenuated delays in larval development due to TOR suppression by rapamycin.

**Figure 4.**
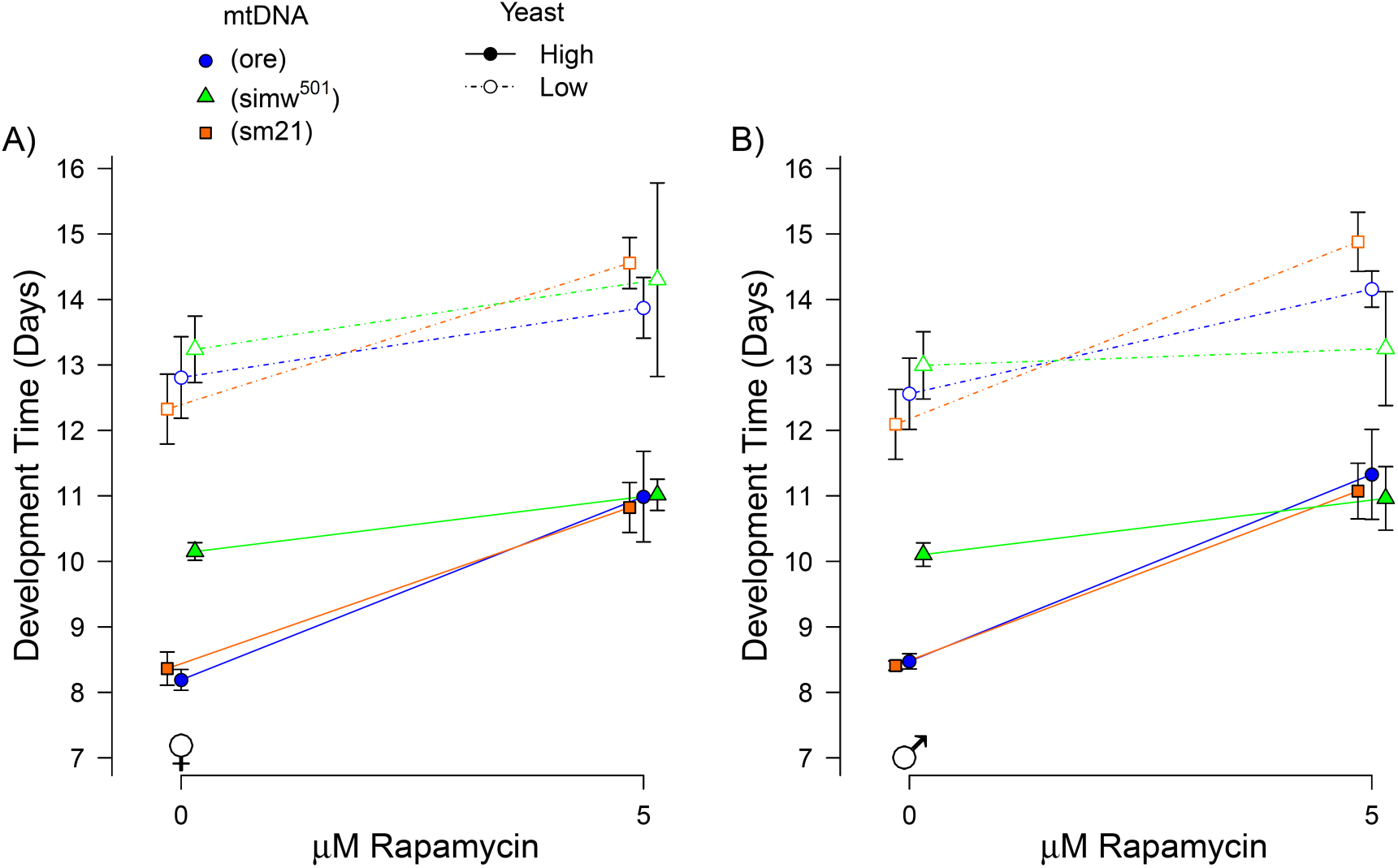
A low-yeast diet attenuated the response of some mitochondrial-nuclear genotypes to a low dose of rapamycin. Similar to (*simw^501^*);*OreR* on a high-yeast diet, the (*ore*);*OreR* genotype had an attenuated response to rapamycin when fed a low-yeast diet, an effect that was somewhat stronger in females (A) than in males (B). Points are average trait values across five replicate vials with 95% CI.

### Females with compromised energy metabolism have decreased immune function

We measured the survival of (*simw^501^*);*OreR* and the compatible mito-nuclear genotypic controls that were infected as adults with the natural *Drosophila* pathogen *P. rettgeri*, as well as flies that were given a sham infection. Survival following infection was significantly affected by mito-nuclear genotype (mtDNA*nuclear*infection: *χ^2^*_2,104_ = 8.51, *P* = 0.014), due primarily to decreased survival of (*simw^501^*);*OreR* females (females, mtDNA*nuclear*infection: *χ^2^*_2,47_ =5.46, *P* = 0.065) (Fig. 5B), as males showed no significant genotype-specific survival differences (males, mtDNA*nuclear*infection: *χ^2^*_2,50_ = 2.9, *P* = 0.23) (Fig. 5A). Consistent with prior studies, we found that most death occurred in the 3-4 days following infection with this pathogen (Duneau, Ferdy, et al. 2017).

**Figure 5.**
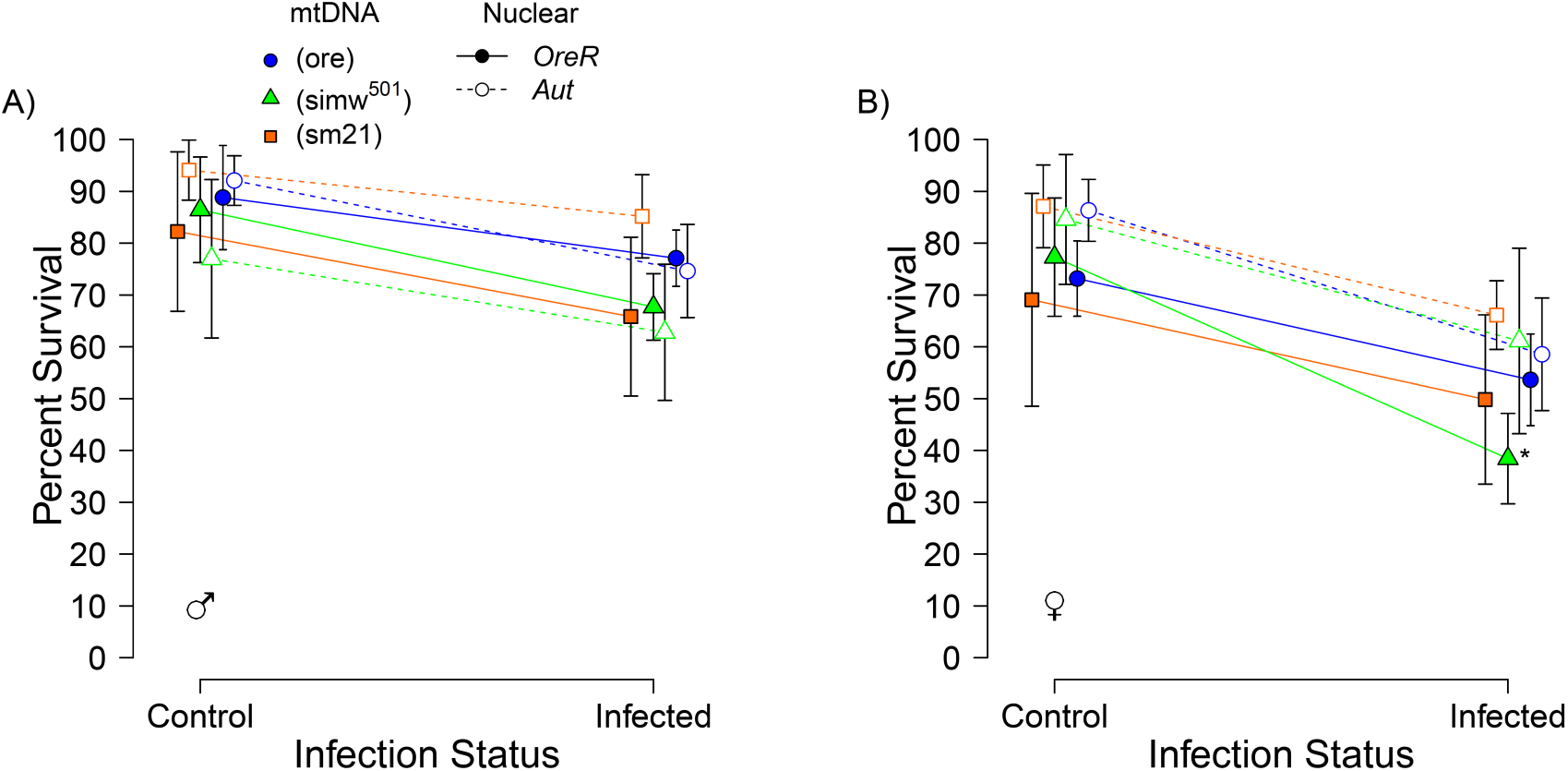
An energetically compromised genotype (*simw^501^*);*OreR* had decreased immune function in females. A) (*simw^501^*);*OreR* females had a greater decrease in survival following infection with the natural pathogen *Providencia rettgeri*, relative to control genotypes. B) Males of this genotype had rates of post-infection survival that were similar to genetic controls. Control infection status refers to sham infections. Points are averages across 5-6 replicate vials with 95% CI.

### Compromised energy metabolism reveals an immunity-fecundity tradeoff

We measured the fecundity of females that survived for five days following either bacterial or sham infection. In control genotypes, there was no evidence for a tradeoff between immunity and fecundity; over the course of three days, females produced similar numbers of offspring whether they had survived a sham infection or a pathogen infection (infection: *F*_1,428_ = 0.0001, *P* = 0.99; mtDNA*nuclear*infection: *F*_1,428_ = 2.7, *P* = 0.10) (Fig. 6, Supplementary Fig. 5). However, (*simw^501^*);*OreR* females that had survived infection with *P. rettgeri* had fewer offspring than sham-infected females of the same genotype (infection: *F*_1,116_ = 3.97, *P* = 0.049) (Fig. 6), an effect that was magnified across the three days of egg production (Supplementary Fig. 5).

**Figure 6.**
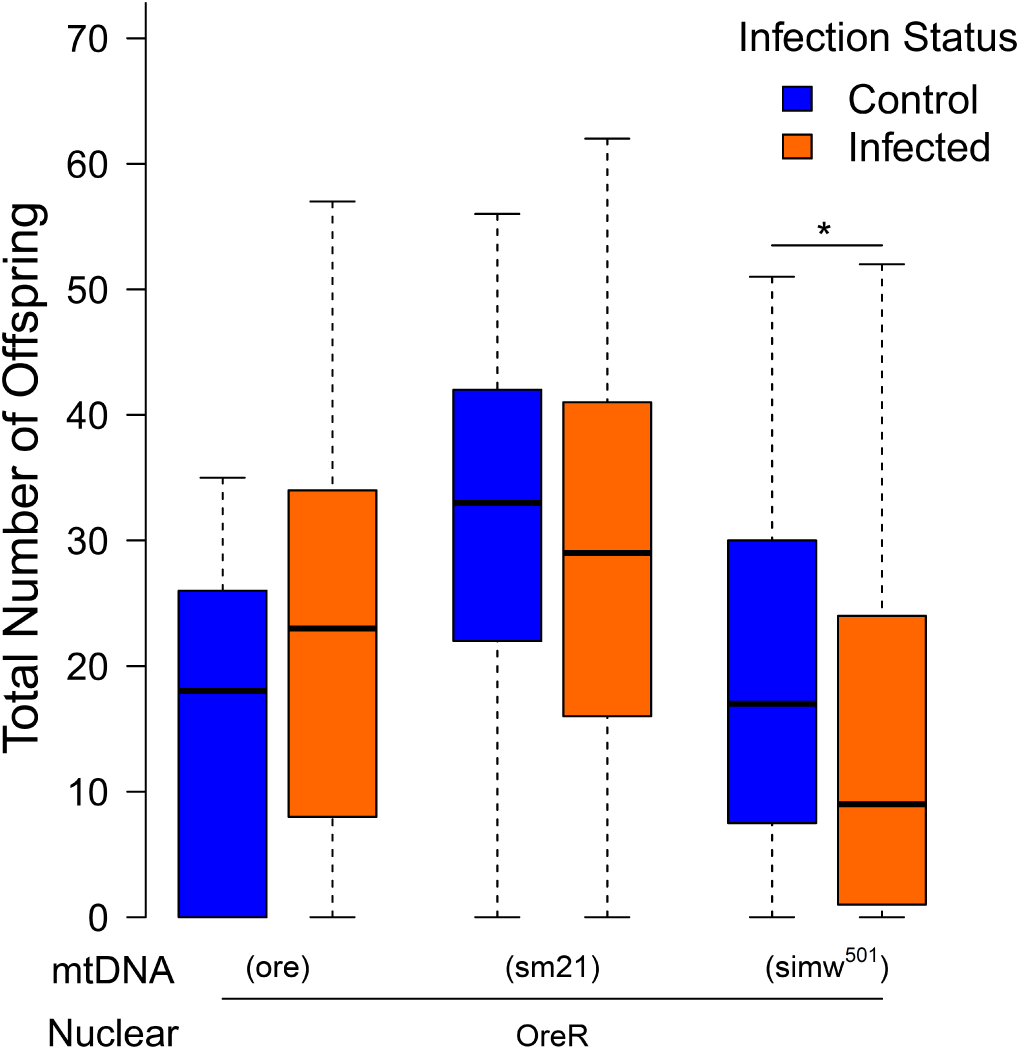
Compromised energy metabolism in (*simw^501^*);*OreR* revealed an immunity-reproduction tradeoff. Surviving infection decreased the total number of offspring produced by (*simw^501^*);*OreR* females, relative to sham-infected females, an effect that was not observed in genotypic controls. The total number of offspring represent those offspring that survived to adult eclosion. Data shown are from the *OreR* nuclear genotype due to the large differences in fecundity between the *OreR* and *Aut* nuclear backgrounds. Data for all genotypes across three days of egg laying are provided in Supplementary Fig. 5. Boxplots represent 15-20 replicate females for each genotype.

## Discussion

Life-history tradeoffs occur due to differential allocation of resources to the competing demands of organismal growth, maintenance, performance, and reproduction (Harshman and Zera 2007; King et al. 2011). These tradeoffs can vary between genotypes within a species or within an individual across life stages (Zera and Larsen 2001), and can be modified by environmental stresses, such as temperature (Partridge et al. 1995), pathogens (Love et al. 2008; McKean et al. 2008; Schwenke et al. 2016; Valtonen and Rantala 2012), and decreased resource availability (Burger et al. 2007). The latter can have particularly strong effects on reproductive fitness that can range from gonadal development (Bergland et al. 2008) to the production of sexual ornaments and signals (Siva-Jothy 2000; Fedorka and Mousseau 2007; Emlen et al. 2012; Gilbert and Uetz 2016; Gilbert et al. 2016). Decreased dietary resources negatively impact ovary development and the number of eggs produced by female *Drosophila* (Drummond-Barbosa and Spradling 2001; Bergland et al. 2008). In other insects, decreased nutritional resources can lead to lower immune activation (Jacot et al. 2005), changes in gene expression related to immune function (Adamo et al. 2016), and decreased fecundity (Stahlschmidt et al. 2013). Remarkably, larvae of the moth *Spodoptera littoralis* prefer diets that maximize the appropriate immune response (Cotter et al. 2011), indicating that energetic-immune interactions are likely important in shaping evolutionary responses to environmental challenges, as well as mediating life-history tradeoffs.

However, nutrient reduction is not always detrimental to immunity (Adamo et al. 2016) or fecundity (May et al. 2015). Short-term starvation can increase infection survival (Brown et al. 2009), and decreased nutrition can increase generalized immune responses, such as phenyloxidase production (Miller and Cotter 2017a) and encapsulation (Saastamoinen and Rantala 2013), despite the fact that immune responses are energetically expensive (Cutrera et al. 2010;Kvidera et al. 2017). It is possible that differences observed between studies are due to differences in the type (generalized vs. specific) of immune response under investigation (Lee 2006), but could also be due to other life-history differences between species (Hawley and Altizer 2011). Our results indicate that genetic variation in mitochondrial and nuclear genomes impacts immune response to a natural bacterial pathogen, and reveals immunity-fecundity tradeoffs in female *Drosophila*, likely due to a compromised mitochondrial ability to convert environmental nutrients to cellular resources.

In response to the natural bacterial pathogen *P. rettgeri, Drosophila* activate the Toll, IMD, and JAK/STAT pathways in the first day of infection and the degree of activation is predictive of survivorship (Sackton et al. 2010; Duneau, Ferdy, et al. 2017). However, natural populations harbor significant genetic variation for surviving infection by *P. rettgeri* and these genetic effects are modified by diet (Howick and Lazzaro 2014). Our results suggest that mutations that impact mitochondrial function may be in important source of genetic variation for immune function in natural populations. Mitochondria have been linked to innate and adaptive immune responses (West et al. 2011; Pourcelot and Arnoult 2014; Weinberg et al. 2015), although mitochondrial genotype does not always decrease post-infection reproduction (Nystrand et al. 2017). While we infer that reduced survival and fecundity in infected (*simw^501^*);*OreR* females is due to a decreased energy supply that cannot meet the demands of immune function, mitochondria also have other roles in immunity that may contribute to this observation, including ROS production, mitochondrial antiviral signaling, and cellular damage responses (West et al. 2011; Pourcelot and Arnoult 2014; Weinberg et al. 2015).

Our data suggest that TOR signaling may be activated and less responsive in energetically inefficient genotypes. External and internal energy sensing is integrated by TOR (Xu et al. 2012; Rider 2016) to regulate growth (Zhang et al. 2000; Kavitha et al. 2014), fecundity (Zhai et al. 2015), and autophagy (Neufeld 2010), and there is some indication of a role for TOR signaling in immunity (Cobbold 2013; Allen et al. 2016). TOR signaling is sensitive to many factors including decreased nutrition (Nagarajan and Grewal 2014), mitochondrial dysfunction (Kemppainen et al. 2016), and overnutrition (Jia et al. 2014), and populations of *Drosophila melanogaster* harbor genetic variation, including mitochondrial, that influences energy sensing via TOR (Villa-Cuesta, Holmbeck et al. 2014; Stanley et al. 2017). Thus, TOR signaling is an important pathway integrating external and internal energetic and immunity status that may influence the evolution of life-history traits in response to the environment. Our results are consistent with other studies that indicate that this pathway may be limited in the extent to which the addition of multiple inputs can continue to cause increased signaling via TOR. Both simulated low nutrition via rapamycin (Villa-Cuesta, Fan, et al. 2014) and genetic activation of TOR (Nagarajan and Grewal 2014) fail to generate the expected phenotypic effects of nutrient limitation. Together, these observations indicate that there may be a threshold for nutrient sensing that, once crossed, prevents further suppression of TOR.

Infection reduced (*simw^501^*);*OreR* survival only in females. In general, male *Drosophila* survive infection better than do females (Short and Lazzaro 2010; Vincent and Sharp 2014; Duneau, Kondolf, et al. 2017), a pattern that we observe here. The relatively higher survival of males could result from sex-specific differences in immune expression due to Y-linked regulation (Fedorka and Kutch 2015), differences in antimicrobial peptide production (Jacobs et al. 2016; Duneau, Kondolf, et al. 2017), or potentially from differential suppression of the immune system by juvenile hormone, which has been shown to underlie differences in immune function between mated and un-mated females (Schwenke and Lazzaro 2017). An energetic explanation may be that females simply have less excess supply to invest in immune function, due to differential costs of gamete production (Bateman 1948; Hayward and Gillooly 2011; McKean et al. 2008; Rolff 2002; Schwenke et al. 2016). Mated females have lower antimicrobial peptide production than non-mated females (Short and Lazzaro 2010), and our results indicate that compromising cellular energy metabolism has greater effects on female immune function and female reproduction, relative to male performance (Hoekstra et al. 2018).

These patterns are counter to the expectation that female *Drosophila* might mount stronger immune responses, because the resulting increase in longevity provides greater lifetime opportunity for reproduction (McKean and Nunney 2005), a pattern that has been observed in many species (Klein 2004; Nunn et al. 2009; Miller and Cotter 2017b). In fact, investment in immunity has been shown to be greater in the sex that has higher investment in offspring, regardless of sex (Roth et al. 2011). However, this pattern may not be observed across all conditions, as environmental effects such as stress can decrease immune responses (Husak et al. 2017). Furthermore, in a study where female *Drosophila* appeared to invest more in immune function than did males, the effects were influenced by the presence of *Wolbachia* (Gupta et al. 2017). While none of our genotypes are infected with *Wolbachia*, understanding the interactions between this endosymbiont and mitochondrial effects on host energetics, immunity and reproduction would provide important insight on the spread of *Wolbachia* in natural populations. An energetic framework that considers how the balance of energy supply and demand differs across external environmental conditions and across internal conditions, such as sex, endosymbiont status and tissue (e.g., ovary versus testes) (Hoekstra et al. 2018), may be a particular useful framework for predicting under what conditions sexes may differ in their immune investment and when genetic variation in mitochondrial function will have sex-specific effects on immune function and tradeoffs between reproduction and immunity (Cressler et al. 2014; Tate and Graham 2015).

## Acknowledgements

We would like to thank Brian Lazzaro for the *Provedencia rettgeri* and for his intellectual contributions to this study. We are grateful for the technical help of Katie Gordon, Rudy Villegas, Abhilesh Dhawanjewar, Cole Julick, and Omera Matoo. We acknowledge the unwavering support of David Rand.

## Authors’ Contributions

JLB, CDM, and KLM conceived and designed the study. JLB and CDM carried out lab work. JLB and KLM drafted in initial version of the manuscript, and all authors revised and gave final approval for publication.

